# Proteomic Characterization of Serum Small Extracellular Vesicles in Human Breast Cancer

**DOI:** 10.1101/2021.11.26.470104

**Authors:** Ganfei Xu, Weiyi Huang, Shaoqian Du, Minjing Huang, Jiacheng Lyu, Fei Zhou, Rongxuan Zhu, Yuan Cao, Jingxuan Xv, Ning Li, Guoying Yu, Binghua Jiang, Olivier Gires, Lei Zhou, Hongwei Zhang, Chen Ding, Hongxia Wang

## Abstract

There is a lack of comprehensive understanding of breast cancer (BC) specific sEVs characteristics and composition on BC unique proteomic information from human samples. Here, we interrogated the proteomic landscape of sEVs in 167 serum samples from patients with BC, benign mammary disease (BD) and from healthy donors (HD). The analysis provides a comprehensive landscape of serum sEVs with totally 9,589 proteins identified, considerably expanding the panel of sEVs markers. Of note, serum BC-sEVs protein signatures were distinct from those of BD and HD, representing stage- and molecular subtype-specific patterns. We constructed specific sEVs protein identifiers that could serve as a liquid biopsy tool for diagnosis and classification of BC from benign mammary disease, molecular subtypes, as well as assessment of lymph node metastasis. We also identified 11 potential survival biomarkers for distant metastasis. This work may provide reference value for the accurate diagnosis and monitoring of BC progression using serum sEVs.

## Introduction

Breast cancer (BC) is one of the most common cancers worldwide and accounts for 30% of female cancers (Kim *et al*, 2012; Liu *et al*, 2021; Siegel *et al*, 2021). A long-term decline in the death rate has been observed since the mid-1970s due to improvements in treatment protocols, including the development of chemotherapy, immunotherapy and targeted therapies. However, improvements in clinical outcomes have slowed over the past decade, and distant metastasis remains the major cause of mortality (Cassetta & Pollard, 2017; Liu *et al*, 2019a; Siegel *et al*., 2021; Yin *et al*, 2014; Zhu *et al*, 2019). The early detection and dynamic assessment of the metastatic status of BC patients are of great value for the treatment and longitudinal analysis of cancer evolution in response to therapy. To achieve this, liquid biopsies utilizing molecular classifiers detected in blood from patients, such as circulating tumor cells, circulating free DNA, and exosomes, offer minimal invasiveness, fewer complications, and an increased ability for longitudinal monitoring compared with traditional tumor tissue biopsies (Wan *et al*, 2017; Yoneda *et al*, 2019). More importantly, liquid biopsy is more informative than single locally restricted biopsies, providing unique information about tumor heterogeneity, clonal evolution, and the potential development of premetastatic cancer cells (Hoshino *et al*, 2020).

Circulating small extracellular vesicles (sEVs), such as exosomes or exosome-like vesicles (ELVs), are 30-150 nm in size and carry a restricted set of nucleic acids, lipids, and proteins (Balaj *et al*, 2011; Johnstone *et al*, 1987; Kim *et al*, 2013; Peinado *et al*, 2011; Raposo & Stoorvogel, 2013; Skog *et al*, 2008; Thakur *et al*, 2014; Thery *et al*, 2009; Valadi *et al*, 2007; Wang & Gires, 2019) that contribute to intercellular communication in normal physiology and pathology (Johnstone *et al*., 1987; Maas *et al*, 2017; Skog *et al*., 2008; Yanez-Mo *et al*, 2015). The functional importance of sEVs has been intensively studied in multiple human cancers, including BC (Hoshino *et al*., 2020). Increasing evidence suggests that sEVs are actively released from cancer cells and markedly affect the tumor microenvironment (TME) as well as the immune ecosystem (Huber *et al*, 2005), thereby constructing distant metastatic niches and facilitating cancer growth (Fang *et al*, 2018; Kralj-Iglic, 2012; Ozer *et al*, 2020) and metastasis (Chen *et al*, 2018; Costa-Silva *et al*, 2015; Hoshino *et al*, 2015; Peinado *et al*, 2012; Zhang & Wang, 2015). Of note, the membrane encapsulation of sEVs promotes their structural integrity, and cargos located within sEVs are more stable than other serological proteins since they have protection against degradation by circulating proteases and other enzymes (Li *et al*, 2017a). Considering their facilitated retrieval and their relatively ubiquitous presence and abundance in serum, sEVs can provide ample materials for downstream analysis in BC detection, prognosis, and therapeutic monitoring as a promising, noninvasive liquid biopsy approach (Choi *et al*, 2021; Lee *et al*, 2018; Li *et al*, 2017b; Wang *et al*, 2018). For instance, Peinado et al. showed that an “sEV protein signature” could identify melanoma patients at risk for metastasis to nonspecific distant sites (Peinado *et al*., 2012). Hoshino et al. identified a specific repertoire of integrins expressed on cancer-derived sEVs, which were distinct from cancer cells, that dictated exosome adhesion to specific cell types and ECM molecules in particular organs (Hoshino *et al*., 2015).

The sEV proteome has been proposed to offer unique advantages as an informative readout for the detection and stratification of BC (Rontogianni *et al*, 2019). Nonetheless, the challenge is to optimize a proteomic profiling approach for sEVs to define and standardize reliable methods. Despite the availability of several public sEV protein databases (*e.g.,* Vesiclepedia (www.microvesicles.org/) (Kalra *et al*, 2012), EVpedia (www.evpedia.info) (Kim *et al*., 2012) and ExoCarta (www.exocarta.org) (Kim *et al*., 2013)), much remains unknown about the sEV proteomes of BC. This includes the definition of (1) markers to distinguish BC from benign disease and healthy state, (2) markers to distinguish diverse molecular subtypes of invasive breast cancer (IBC), (3) markers to predict lymph node (LN) metastases, and (4) the open question of whether molecules present on IBC-derived sEVs are “addressing” them to specific organs. These unresolved problems highlight the need for a better understanding of the protein composition of BC-derived sEVs that could qualify them as biomarkers for clinical application, with a specificity and sensitivity mostly superior to those of traditional serum markers. To address these aims, mass spectrometry-based proteomic profiling is emerging as a strategy to gain insight into the biological cargos, functions, and clinical potential of sEVs (Wang *et al*, 2020).

Here, we applied a mass spectrometry-based, data-independent acquisition (DIA) quantitative approach to determine the proteomic features of human serum sEVs derived from patients with BC, benign mammary disease (BD), and healthy donors (HDs). In total, we identified 9,589 proteins from 167 analyzed samples with a mean of 1,695 proteins quantified per sEV sample. Classification of the pathways related to the enriched proteins revealed that proteins preferentially packaged in BC-sEVs correlated with interferon γ-mediated signaling as well as pathways associated with immune response regulation, antigen processing and presentation, glycolysis and angiogenesis. By examining the sEV proteomes, we constructed specific sEV protein identifiers that could serve as a liquid biopsy tool for the diagnosis and classification of BC from BD and its molecular subtypes, as well as the assessment of LN metastasis. Of note, we found that adipocytes play an important role in the LN metastasis of BC. We also identified 11 potential survival markers for distant BC metastasis and 2 potential survival markers for lung metastasis. This work may provide reference value for the accurate diagnosis and monitoring of BC progression using serum sEVs, and the identification of novel molecules packaged in sEVs offers an opportunity for the targeted therapy of BC in the future.

## Results

### Proteomic characterization of BC-derived sEVs

To elucidate the proteomic profile of BC-derived sEVs, we purified sEVs from 167 human serum samples derived from BC patients (n = 126), BD patients (n = 17), and HDs (n = 24) by differential ultracentrifugation as described in the Methods and in accordance with previously reported protocols (Colombo *et al*, 2014; Peinado *et al*., 2012; Xu *et al*, 2016) (Fig 1A and C). All samples were collected prospectively from treatment-naive stage I-IV BC patients (Fig 1B, Appendix Table 1). Under transmission electron microscopy (TEM) in combination with nanoparticle tracking analysis (NTA), the isolated sEVs appeared as morphologically uniform vesicular structures 30-150 nm in size surrounded by a double-layer membrane (Fig 1D, Appendix Fig S1A). sEV samples were verified by immunoblotting analyses using the conventional markers CD9, CD63, TSG101, and ALIX, while we examined 24 sEV markers in our proteomics data (Hoshino *et al*., 2020) (Fig 1E, Appendix Fig S1B). Clinical data, including sex, age at diagnosis, tumor staging, BC subtypes, LN status, distant metastasis, and survival, are summarized in Fig 1B and Table S1.

**Figure 1.**
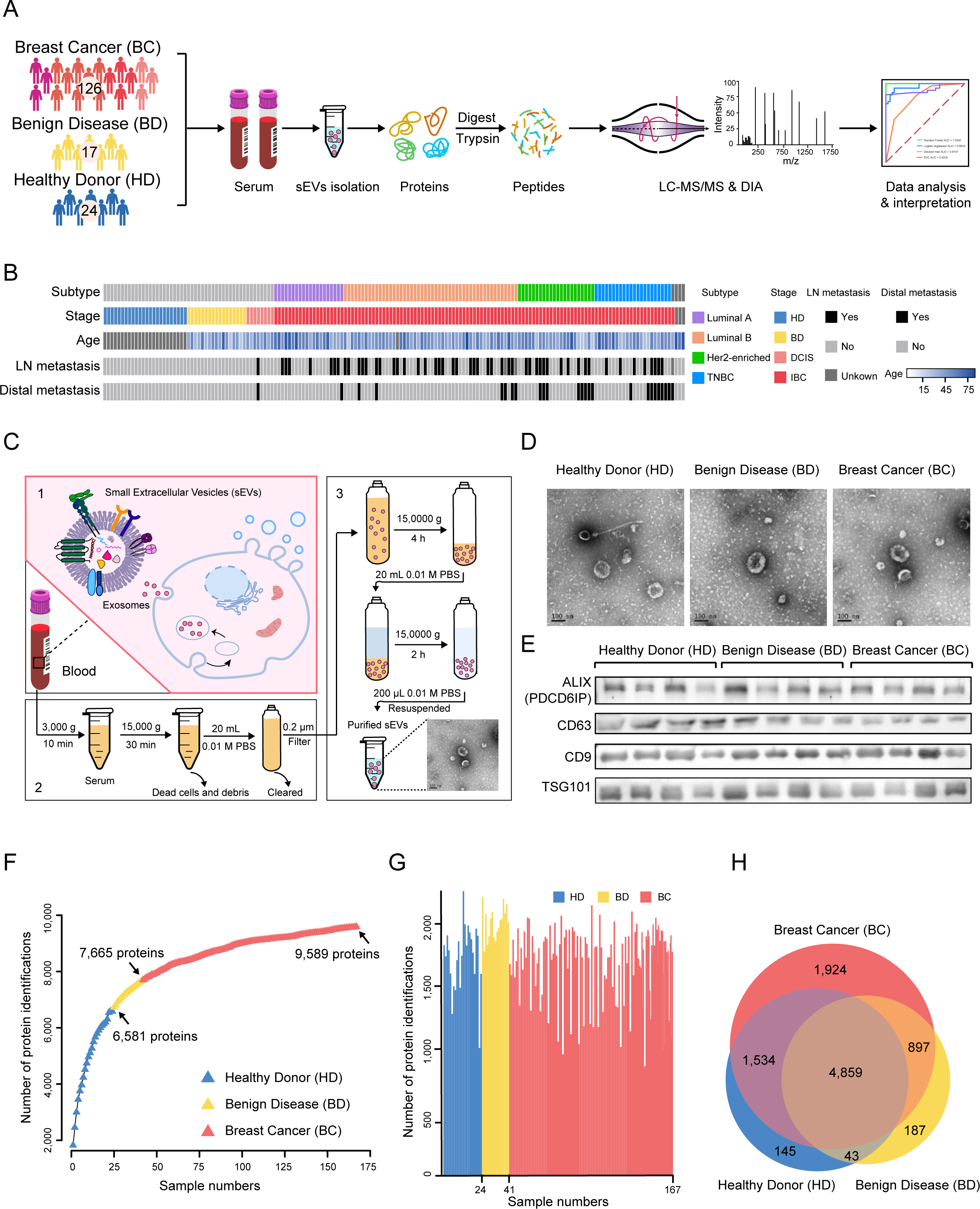
Overview of the proteomic characterization of breast cancer sEVs. A Overview of the experimental design and the number of samples for proteomics analyses. B Clinical parameters are indicated in the heatmap. C Schematic diagram of the extraction process of serum-derived sEVs. D Representative TEM images of purified EVs. Scale bar–100 nm. E Immunoblots showing the expression levels of ALIX (PDCD6IP), CD63, CD9, and TSG101 in the purified EVs. F Cumulative number of protein identifications. Red denotes BC samples (n = 126), yellow denotes BD samples (n = 17), and blue denotes HD samples (n = 24). G The number of proteins identified in 167 samples. Red denotes BC samples (n = 126), yellow denotes BD samples (n = 17), and blue denotes HD samples (n = 24). H Venn diagram depicting the numbers of proteins detected in BC-, BD-, and HD-sEVs.

A proteomic database of serum sEVs was constructed using label-free LC-MS/MS analysis, identifying 9,589 proteins in total from the 167 analyzed samples at a protein- and peptide-level FDR of less than 5% (Fig 1F). The protein abundance was first calculated by iBAQ and then normalized as FOT, allowing for comparison among different experiments. The mean number of proteins detected per sEV sample was 1,695 (range 793 to 2,253 proteins) (Fig 1G). In general, 1,924, 187, and 145 unique sEV proteins were identified in BC, BD, and HD samples, respectively (Fig 1H). Globally, the dynamic range of proteins detected spanned eight orders of magnitude (Appendix Fig S1C). Collectively, these data were consistent with previous reports that sEV protein profiles differ significantly depending on the sample source (Wu *et al*, 2019), and sEVs released by BC cells and from other cancer cells may carry more encapsulated cargos for signal transfer to induce the malignant transformation and proliferation of recipient cells (Milane *et al*, 2015).

### BC-derived sEVs exhibited specific signatures related to immune response, metabolism, and metastasis

Next, proteomic data were analyzed to determine the characteristics of BC-derived sEVs. PCA demonstrated a clear distinction among the three different types of samples, which further highlighted the diverse proteomic patterns among BC-, BD-, and HD-sEVs that underpinned our stratification analysis (Appendix Fig S2A).

To decipher the protein network associated with BC tumorigenesis, we identified 287, 602, and 112 proteins that were significantly overrepresented in the BC (BC_mean_/BD_mean_ > 2-fold and BC_mean_/HD_mean_ > 2-fold), BD (BD_mean_/BC_mean_ > 2-fold and BD_mean_/HD_mean_ > 2-fold) and HD (HD_mean_/BC_mean_ > 2-fold and HD_mean_/BD_mean_ > 2-fold) samples, respectively (see Materials and Methods). Clustering and cluster-specific enrichment analyses of these proteins using GOBP and Reactome pathway annotations showed that these differentially enriched proteins were involved in distinctive biological processes and pathways (Fig 2A, Appendix Table 2). Specifically, COPI-mediated anterograde transport (Fisher’s exact test, *p* = 3.88e-3), vesicle-mediated transport (Fisher’s exact test, *p* = 1.26e-4), and regulation of actin dynamics for phagocytic cup formation-related proteins (Fisher’s exact test, *p* = 2.48e-5) (*i.e.,* ADD2, ARF5, ARPC1A, IGHV3-53, IGHV4-39, SSC5D, and COPE) were enriched in HD samples (Fig 2A and B, Appendix Table 2). BD-sEVs were characterized by proteins related to cell-cell adhesion (Fisher’s exact test, *p* = 2.98e-19) (*i.e.,* STAT1, PTPN1, RPL24, and FNBP1L), cholesterol metabolic process (Fisher’s exact test, *p* = 3.34e-7) (*i.e.,* PON1, APOC1, APOA2, ANGPTL3, and LIPC), and response to estrogen (Fisher’s exact test, *p* = 2.13e-2) (*i.e.,* F7, LDHA, HSP90AA1, IGFBP2, and CTNNA1) (Fig 2A and B, Appendix Table 2). Of note, BC-sEVs exhibited specific signatures related to the immune response, metabolism, and metastasis, potentially reflecting the functional roles and molecular heterogeneity of sEVs during BC tumorigenesis and progression. Classification of the pathways related to the enriched proteins from BC-sEVs revealed that these selectively packaged proteins are involved in the interferon γ-mediated signaling pathway (Fisher’s exact test, *p* = 4.94e-4) (*i.e.,* HCK, HLA-H, HLA-B, HLA-C, HLA-A, HLA-G, and CD44), regulation of immune response (Fisher’s exact test, *p* = 4.61e-5) (*i.e.,* IGLV3-25, COL3A1, CXADR, IGLV3-27, HLA-A, IGLV7-43, and PVR), antigen processing and presentation (Fisher’s exact test, *p* = 1.16e-5) (*i.e.,* ITGB1, IGLV3-25, CXADR, IGLV3-27, IGLV7-43, PVR, and HLA-G), glycolytic process (Fisher’s exact test, *p* = 1.29e-3) (*i.e.,* GPI, PGK1, PGAM4, PGK2, and PGM1), and angiogenesis (Fisher’s exact test, *p* = 3.92e-2) (*i.e.*, GPI, RNF213, ANGPTL6, MMP2, PECAM1, CYP1B1, NAA15, and TYMP) (FC > 2, one-way ANOVA *p* < 0.05) (Fig 2A and B, Appendix Table 2). Notably, in the Tang et al. BC cohort (Tang *et al*, 2018), among sEV proteins that were specifically highly expressed in BC samples, patients with high expression of MMP2 and TYMP appeared to have poor prognostic outcomes (log rank test, *p* < 0.05) (Fig 2C). These findings that BC-, BD-, and HD-sEV cargos are distinct and related to singular cellular processes suggest that sEV protein packaging into sEVs is heterogeneous and reflects BC biology.

**Figure 2.**
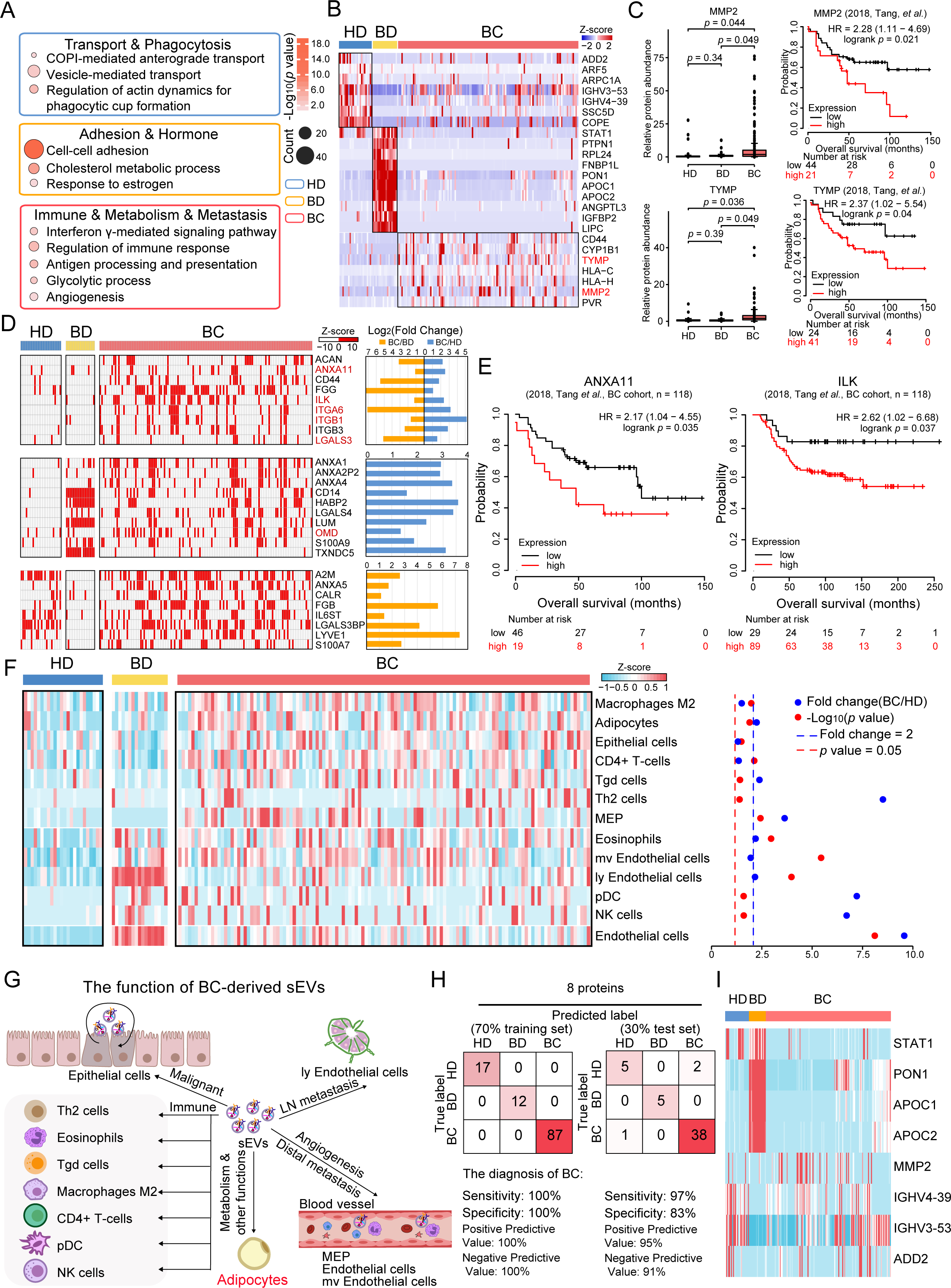
Proteomics features of BC-, BD- and HD-derived sEVs. A The bubble plot indicates the distinctive biological pathways of BC, BD, and HD. Red box, BC; yellow box, BD; blue box, HD. See Table S2. B Differentially expressed proteins in the distinctive biological pathways of BC, BD, and HD. Fold change > 2 and one-way ANOVA *p* < 0.05. C Two proteins (MMP2 and TYMP) differentially expressed in BC, BD, and HD (*p* value from Student’s *t* test) and their association with clinical outcomes in BC (2018, Tang et al., BC cohort, n = 118) (*p* value from log rank test). D sEV DAMP molecules were enriched in BC and found in > 50% of BC samples, with > 2-fold difference and one-way ANOVA *p* < 0.05. E DAMP molecules enriched in BC-sEVs were significantly associated with clinical outcomes in BC (2018, Tang et al., BC cohort, n = 118) (*p* value from log rank test). F Distinctive tumour microenvironment in BC. See Table S2. G Functions of BC-derived sEVs. These sEVs impact the tumour microenvironment by promoting tumour cell growth and progression, modulating immune responses, regulating angiogenesis and inducing metastatic behaviour through MEPs, endothelial cells, and Mv endothelial cells. H Classification error matrix of the training set (70%) and test set (30%) for the 8 proteins using the random forest classifier. The number of samples identified is noted in each box. I Proteins with the highest predictive values in classifying BC, BD and HD samples by XGBoost.

### Specific damage-associated molecular pattern (DAMP) molecules are packaged in BC-derived sEVs

Recent advances have indicated that DAMP molecules, such as nucleic acids, histones, high mobility group box 1, S100, and heat shock proteins, act as endogenous ligands of innate immune receptors and are linked to the immune response and cancer progression (Becker *et al*, 2016). In total, we identified 210 different DAMPs in all sEV datasets (Appendix Fig S2B, Appendix Table 2). Specifically, the analysis identified 197, 145, and 157 DAMPs in BC-, BD-, and HD-sEVs, respectively, suggesting that more DAMPs were enriched in BC samples than in BD and HD samples (Appendix Fig S2B). Thirty-two of these DAMPs were identified only in BC-sEVs, 9 DAMPs only in BD-sEVs, and 4 DAMPs only in HD-sEVs (Appendix Fig S2B). Of all DAMPs identified in BC-sEVs, 27 DAMPs (*e.g.,* ACAN, ANXA11, and CD44) were shared by > 50% of BC samples and were enriched compared to BD-sEVs and/or HD-sEVs (Fig 2D).

Among them, 9 DAMPs, including aggrecan (ACAN), annexin A11 (ANXA11), CD44, fibrinogen gamma chain (FGG), integrin-linked kinase (ILK), LGALS3, and several ITGs (ITGA6, ITGB1, and ITGB3), were exclusively present in BC-sEVs versus BD- and HD-sEVs, suggesting that they are specific sEV markers in BC development and progression (Fig 2D). ITGA6, ITGB1, and ITGB3 are members of the integrin family of proteins involved in cell adhesion and recognition in a variety of processes, including tissue repair, hemostasis, immune response, and metastatic dissemination of cancer cells (Laudato *et al*, 2017; Wang *et al*, 2019b). ANXA11 and LGALS3 are associated with the progression of some cancers (Liu *et al*, 2019b; Wang *et al*, 2019a). Another 10 DAMP proteins were highly enriched in both BC- and BD-sEVs: ANXA1, ANXA2P2, ANXA4, CD14, HABP2, LGALS4, LUM, OMD, S100A9, and TXNDC5, whereas they were rarely detected in HD samples, suggesting that they represent sEV DAMPs shared across BC and BD (Fig 2E). Interestingly, our analyses revealed that 8 DAMP molecules (A2M, ANXA5, CALR, FGB, IL6ST, LGALS3BP, LYVE1, and S100A7) were abundantly expressed in both BC- and HD-sEVs (Fig 2E). This finding is consistent with previous studies reporting that the noncancer-derived sEV proteome is as informative as the cancer-derived sEV proteome in specific cancer types (Hoshino *et al*., 2020). It is worth noting that 6 of these molecules (ANXA11, ILK, ITGA6, ITGB1, LGALS3, and OMD) were highly expressed in BC and were associated with poor prognosis in the Tang et al. BC cohort (Tang *et al*., 2018) (Fig 2E, Appendix Fig S2C).

### Possible intercellular communication network diagram of BC-driven sEVs in the TME

Previous evidence suggests that sEVs interact with recipient immune cells to participate in TME remodeling, an effect that is mediated by encapsulated molecular cargos derived from parent cancer cells (Becker *et al*., 2016). Thus, the proteomics profile of BC-sEVs may reflect the status of corresponding immune cells in the TME. To further map the differentially enriched sEV proteins to the immune response, we performed cell type deconvolution analysis using xCell (Aran *et al*, 2017). A heatmap of overall and type-specific enrichment scores was constructed to identify the immune landscape of BC (Fig 2F). Specifically, the enrichment scores of macrophages M2, adipocytes, epithelial cells, CD4+ T cells, γδ T cells (Tgd), Th2 cells and megakaryocyte-erythroid progenitor cells (MEPs) were significantly elevated in BC-sEVs compared to HD-sEVs, with FC > 1.3 and Student’s *t* test *p* value < 0.05 (Fig 2F, Appendix Table 2). The analysis suggested a possible intercellular communication network of BC-driven sEVs in the TME when we inferred the relative abundance of various immune cell subtypes in the TME. MEPs represent a bipotent transitional state that is permissive to the generation of unipotent progenitors of megakaryocytic or erythroid lineages (Xavier-Ferrucio *et al*, 2019). Adipocytes in the TME play dynamic and sophisticated roles in facilitating BC development (Cao, 2019). These BC-derived sEVs may impact the TME by promoting tumor cell growth and progression, modulating immune responses, regulating angiogenesis and inducing metastatic behavior through MEPs, endothelial cells, and mv endothelial cells (Fig 2G).

### Eight-protein diagnostic model to distinguish BC from BD and the healthy population

To further assess whether sEV proteins could be used as a liquid diagnostic tool to discriminate cancers from noncancers, we next sought to determine shared and unique sEV proteins by performing pairwise comparisons of proteomes between BC-, BD-, and HD-sEVs. We applied the XGBoost classifier, which is robust to noise and overfitting, to verify a distinct sEV protein subset that can accurately distinguish the BC, BD and HD samples.

To train and subsequently test the model, sEV samples were evenly partitioned based on the sample source, and 70% of samples were used as a training set, with the remaining 30% used as an independent test set. Applying 5-fold cross-validation to the training set, a combination of 8 sEV proteins (STAT1, PON1, APOC1, APOC2, MMP2, IGHV4-39, IGHV3-53, and ADD2) was used to construct a signature that yielded a sensitivity of 100% and specificity of 100% for discriminating BC from BD and HD (Fig 2H-I, Appendix Fig S2D). Notably, when applying this eight-protein identifier to sEV samples of the independent test set, the model achieved 97% sensitivity and 83% specificity in the diagnosis of BC (Fig 2H).

### Proteomic characteristics of sEVs derived from four clinical subtypes of BC

IBC is a highly heterogeneous disease that can be categorized into various intrinsic or molecular subtypes, which are differentially correlated with clinical presentation, prognosis, distant metastasis, and response to therapy. Molecular subtypes are defined based on the gene expression signature and protein expression of estrogen receptor (ER), progesterone receptor (PR), human epidermal growth factor receptor 2 (Her2), and proliferative cell nuclear antigen (Ki67) (Li *et al*, 2021; Peng *et al*, 2019; Vallejos *et al*, 2010). We reasoned that since the biological behavior of IBC cells differs significantly among IBC subtypes, biological cargos carried by sEVs may vary among diverse molecular subtypes. To distinguish proteomic landscapes among diverse molecular subtypes of IBC and identify drivers that boost intertumoral heterogeneity and cancer evolution, we analyzed sEV samples from luminal A (ER+/PR+, low-grade and low-Ki67 index, n = 20), luminal B (ER+/PR+ of higher grade and proliferative index, n = 50), Her2-enriched (Her2+ with or without ER, n = 21), and triple-negative (ER-PR-Her2-, TNBC, n = 23) IBCs in our cohort. PCA demonstrated a clear distinction among the different molecular subtypes, which further highlighted the distinct proteomic patterns among several clinical subtypes of IBC samples (Appendix Fig S3A).

Next, we applied a *t* test with a nominal *p* value cut-off of < 0.05 and identified 87, 82, 83, and 104 sEV proteins that were significantly overrepresented in luminal A (FC (luminal A/any of the other three subtypes) > 2), luminal B (FC (luminal B/ any of the other three subtypes) > 2), Her2-enriched (FC (Her2-enriched/ any of the other three subtypes) > 2), and TNBC (FC (TNBC/ any of the other three subtypes) > 2) samples (see Materials and Methods). Clustering and cluster-specific enrichment analyses of the enriched proteins using GOBP and KEGG pathway annotations showed the distinctive biological processes and pathways represented in luminal A, luminal B, Her2-enriched, and TNBC samples (Fig 3A and B, Appendix Table 3). Specifically, luminal A-derived sEVs were characterized by proteolysis involved in cellular protein catabolic processes (*i.e.,* PSMB7, PSMB2, FAP, and CAPN2) (Fisher’s exact test, *p* = 1.23e-3) and positive regulation of protein insertion into mitochondrial membrane involved in apoptotic signaling pathway (*i.e.,* YWHAB, YWHAG, and YWHAH) (Fisher’s exact test, *p* = 7.69e-3). Luminal B-derived sEVs were characterized by cellular response to insulin stimulus (*i.e.,* RAB10, PKLR, GOT1, and STAT1) (Fisher’s exact test, *p* = 4.41e-3) and response to hypoxia (*i.e.,* ALAD, VCAM1, PKLR, and HSPD1) (Fisher’s exact test, *p* = 3.77e-2). Her2-enriched sEV-enriched proteins were related to cellular response to reactive oxygen species (*i.e.,* PRDX1, TXN, and SOD3) (Fisher’s exact test, *p* = 1.34e-2), glucose metabolic process (*i.e.,* FABP5, GAA, BPGM, and GAPDH) (Fisher’s exact test, *p* = 3.47e-3), and keratinization (*i.e.,* CASP14, KRT17, and TGM3) (Fisher’s exact test, *p* = 1.99e-2). TNBC samples were characterized by platelet degranulation (*i.e.,* AHSG, ACTN4, PPBP, TLN1, and PF4) (Fisher’s exact test, *p* = 2.67e-3), blood coagulation (*i.e.,* EHD1, COL1A1, PROC, COL1A2, F11, and PRKACB) (Fisher’s exact test, *p* = 3.74e-3), adaptive immune response (*i.e.,* DBNL, ANXA1, ERAP2, and ICOSLG) (Fisher’s exact test, *p* = 5.03e-2), and platelet activation (*i.e.,* COL1A1, COL1A2, SAA1, and PF4) (Fisher’s exact test, *p* = 2.67e-2) (Fig 3B and C, Appendix Table 3).

**Figure 3.**
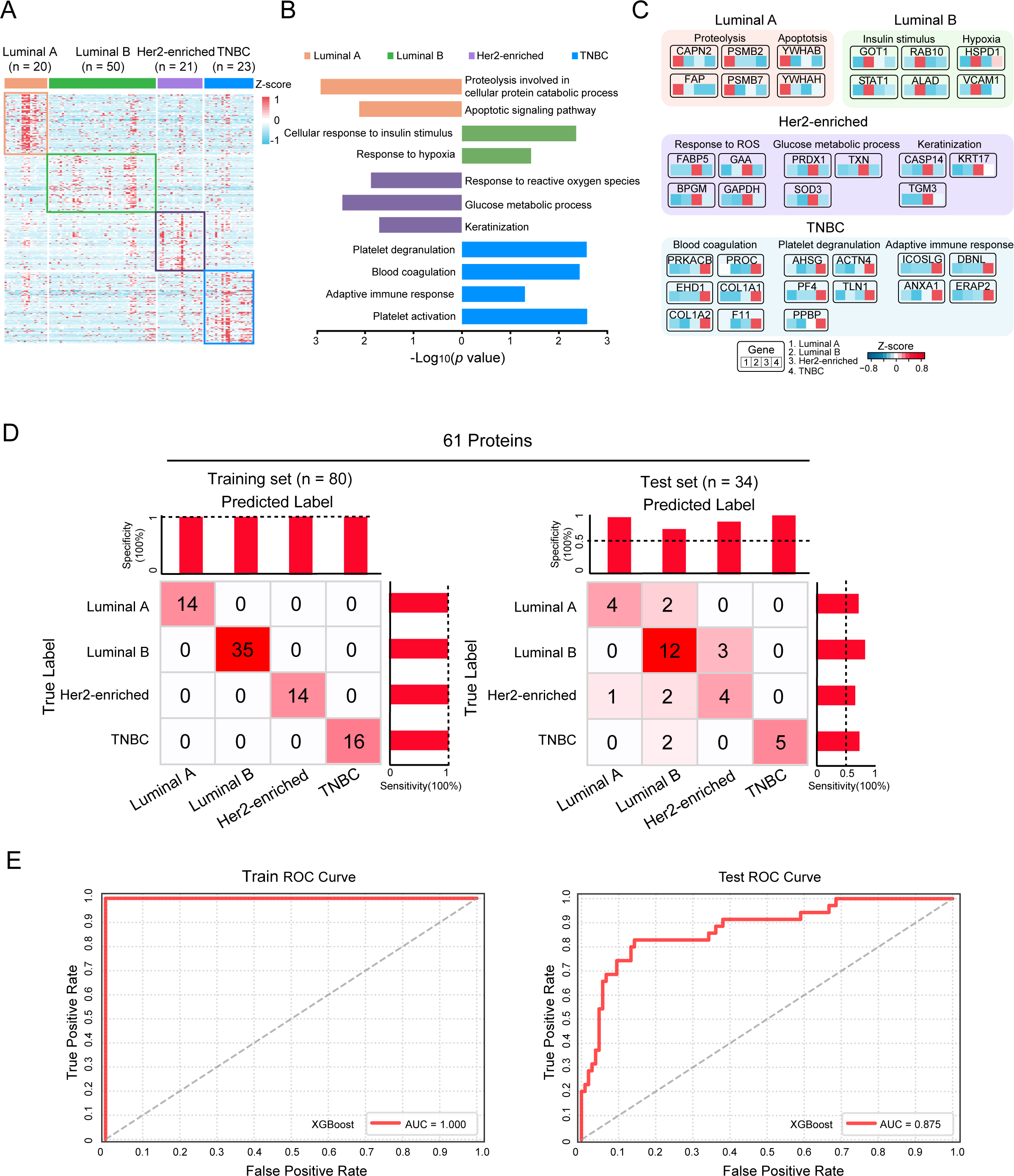
Proteomic landscapes of four clinical subtypes of BC-derived sEVs. A Differentially expressed proteins in luminal A, luminal B, Her2-enriched, and TNBC samples and found in > 50% of the corresponding samples, with > 2-fold difference from the other three subtypes. B Gene Ontology biological processes (GOBPs) revealed pathways that were significantly enriched in luminal A, luminal B, Her2-enriched and TNBC samples (Fisher’s exact test, *p* < 0.05). See Table S3. C Differentially expressed proteins in luminal A, luminal B, Her2-enriched, and TNBC samples. See Table S3. D Classification error matrix of the training set (70%) and test set (30%) for the 61 proteins using the XGBoost classifier. The number of samples identified is noted in each box. The bar chart above represents the predictive specificity of each subtype. The bar chart on the right represents the predictive sensitivity of each subtype.

Collectively, these data suggested that proteomic profiles of serum-derived sEVs reflect selective packaging, which represents an informative readout and differs among diverse subtypes of BCs.

### sEV-based classifier discriminates BC subtypes

To further investigate the clinical significance of the differentially enriched protein cargos, we addressed whether they could be utilized as a novel liquid biopsy method to distinguish diverse clinical subtypes in clinical practice. Employing XGBoost classification, which is robust to noise and overfitting, we constructed a 61-protein classifier model that can accurately discriminate the luminal A, luminal B, Her2-enriched, and TNBC subtypes. To train and subsequently test the model, 70% of samples were used as a training set, with the remaining 30% used as an independent test set, in the same manner as previously described. Similar to our analysis of BC versus non-BC-sEVs, we constructed a 61-protein classifier model using the XGBoost classifier. To test the 61-sEV protein model, 5-fold cross-validation of the training set was performed and yielded a sensitivity of 100% and a specificity of 100% for each molecular subtype (Fig 3D, Appendix Fig S3B). When applying the 61-protein classifier to the independent test set, the model achieved 67% sensitivity and 97% specificity in the diagnosis of luminal A, 80% sensitivity and 70% specificity in diagnosis of luminal B, 57% sensitivity and 89% specificity in diagnosis of Her2-enriched, and 71% sensitivity and 100% specificity in diagnosis of TNBC (Fig 3D, Appendix Fig S3B). The receiver operating characteristic (ROC) curve derived from the 61-protein signature showed good sensitivity and specificity, with an area under the curve (AUC) of 1.0 (Fig 3E). Then, the 61-protein signature was validated in the test set, resulting in a ROC curve with an AUC of 0.875 (Fig 3E).

Thus, serum sEV proteomes can be beneficial in determining the BC subtype for dynamic monitoring in patients during tumor progression, avoiding repeated tissue biopsies.

### Adipocytes play an important role in LN metastasis of BC

Furthermore, to elucidate the mechanism of LN metastasis in IBC, we analyzed sEV proteins of IBC patients with LN metastases (IBC_LN, n = 51) and without LN metastases (IBC_Pure, n = 54). PCA clearly distinguished between IBC_LN and IBC_Pure samples at the protein level, which further highlighted the diverse proteomic patterns between sEVs from IBC_LN and IBC_Pure samples (Appendix Fig S4A). We applied Student’s *t* test with a nominal *p* value cut-off of < 0.05 and identified significantly enriched sEV-derived proteins in IBC_LN compared with IBC_Pure (FC > 2). The results are summarized in the volcano plot shown in Fig S4B, and the most prominent proteins are indicated (Appendix Fig S4B).

We further performed clustering and cluster-specific enrichment analyses of the upregulated proteins using gene set enrichment analysis (GSEA). We found that IBC_LN samples were characterized by proteins related to hallmarks of adipogenesis (Fig 4A). To investigate the immune landscapes of the IBC_Pure and IBC_LN groups, the abundance of 16 different cell types was computed using xCell based on proteomic data of sEVs retrieved from the blood of the 105 abovementioned IBC samples (Fig 4B, Appendix Table 4). We found that the enrichment scores of B cells, basophils, CD4+ T cells, CD4+ naive T cells, dendritic cells (DCs), mesangial cells, activated dendritic cells (aDCs), and immature dendritic cells (iDCs) were higher in the IBC_Pure group than in the IBC_LN group. On the other hand, enrichment scores for adipocytes, CD8+ T cells, CD8+ naive T cells, multipotent progenitors (MPPs), macrophages, megakaryocytes, platelets, and sebocytes were higher in the IBC_LN group than in the IBC_Pure group (FC > 1.5, Student’s *t* test *p* < 0.05) (Fig 4B, Appendix Table 4). The enhanced adipocyte enrichment scores in sEVs from IBC_LN samples attracted our attention (Fig 4B, Appendix Fig S4C, Table 4). There was a positive correlation between adipogenesis and adipocytes (Spearman rho = 0.188, *p* = 5.507e-02) (Appendix Fig S4D).

**Figure 4.**
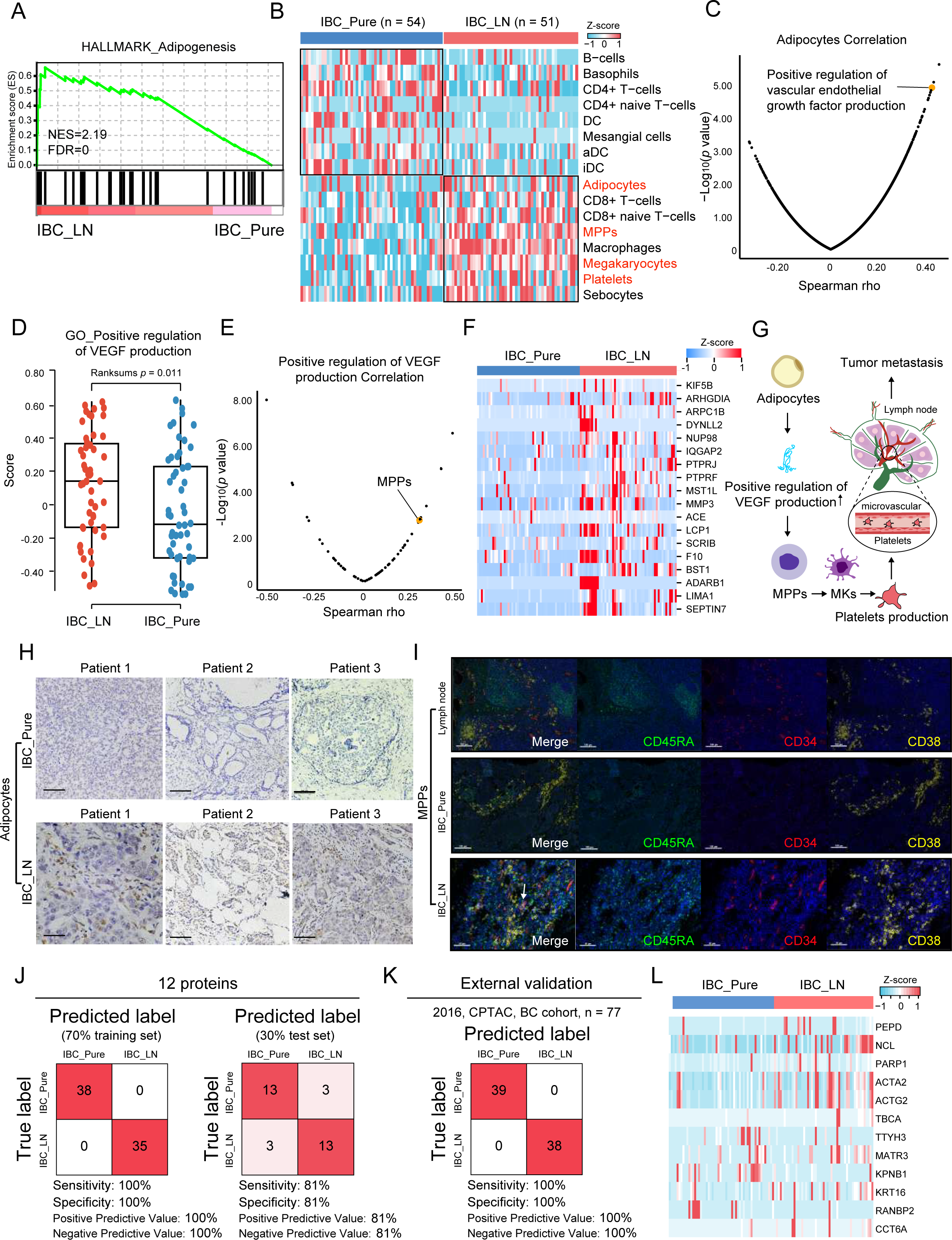
Potential prognostic biomarkers for IBC patients with lymph node metastases. A GSEA of the proteomic data of 105 breast cancer samples revealed that adipogenesis was significantly upregulated in IBC_LN. IBC_LN: IBC patients with lymph node metastases. B Distinctive tumour microenvironment between IBC_Pure and IBC_LN. IBC_Pure: IBC patients without lymph node metastases; IBC_LN: IBC patients with lymph node metastases. See Table S4. C Correlation between adipocytes and the pathway of positive regulation of VEGF production. Spearman rho = 0.412, Wilcoxon rank sum test, *p* = 1.242e-05. D Comparison of the scores of positive regulation of VEGF production between IBC_LN and IBC_Pure. The *p* value was calculated by the Wilcoxon rank sum test. The line and box represent median and upper and lower quartiles, respectively. E Correlation between the pathway of positive regulation of VEGF production and MPPs. Correlation coefficients and *p* values were calculated by the Spearman correlation method. F Molecules highly associated with platelets were expressed in IBC_Pure and IBC_LN. G The pattern diagram shows the process by which adipocytes activate MPPs to generate MEPs and MKs through positive regulation of VEGF production and finally produce platelets. The produced platelets helped breast cancer cells migrate to the lymph nodes. H Representative immunohistochemical images of adipocytes labelled with PPRGg. Images revealed that adipocytes prolifically grew in lymph node metastases of BC compared to primary breast cancer. I Representative fluorescence microscopy images of MPPs labelled with CD45RA (green), CD34 (red), and CD38 (yellow). Images revealed the presence of MPPs in lymph node metastases of BC, which were rare in normal lymph nodes and primary breast cancer. J Classification error matrix of the training set (70%) and test set (30%) for the 12 proteins using the XGBoost classifier. The number of samples identified is noted in each box. K Classification error matrix of the external validation set (2016, CPTAC, BC cohort, n = 77) for the 12 proteins using the XGBoost classifier. The number of samples identified is noted in each box. L Proteins with the highest predictive values in classifying IBC_Pure and IBC_LN samples by XGBoost.

Adipocytes were correlated with the VEGF signaling pathway (Fig 4C), and the VEGF signaling pathway was upregulated in the IBC_LN group (Fig 4D). A previously reported comparative cytokine array analysis of adipocyte-conditioned medium (ACM) revealed the upregulation of a group of cytokines belonging to the VEGF signaling pathway in ACM (Sahoo *et al*, 2018).

The VEGF signaling pathway was correlated with MPPs (Fig 4E), which were upregulated in the IBC_LN group (Appendix Fig S4E) and positively correlated with the coagulation pathway (Spearman rho = 0.295, p value = 2.216e-03) (Appendix Fig S4F). At the same time, platelets were positively correlated with the coagulation pathway (Spearman rho = 0.209, p value = 3.225e-02) (Appendix Fig S4G). The enrichment scores of platelets were upregulated in the IBC_LN group (Appendix Fig S4H). Experimental evidence has highlighted platelets as active players in all steps of tumorigenesis, including cancer growth, cancer cell extravasation and metastasis (Haemmerle *et al*, 2018). Many of the molecules that are highly associated with platelets are angiogenesis- and metastasis-related molecules (*e.g.,* KIF5B, ARHGDIA, ARPC1B, DYNLL2, NUP98, IQGAP2, PTPRJ, PTPRF, MST1L, and MMP3) (Fig 4F, Appendix Fig S4I). In addition, we found that adipocytes, MPPs, and MEPs were significantly increased in the tissue samples of 40 additional IBC patients (IBC_Pure, n = 12; IBC_LN, n = 28), and the platelet count in BC patients with LN metastasis (n = 43) was significantly higher than that in BC patients without LN metastasis (n = 45) in our cohort (Student’s *t* test, *p* < 0.05) (Fig 4H and I, Appendix Fig S4J and K).

### Twelve-protein diagnostic model for LN metastasis

To generate a protein signature that stratifies patients with or without LN metastases, we performed random forest classification to identify a subset of sEV proteins that accurately discriminates between IBC_LN and IBC_Pure samples. As before, sEV samples were evenly partitioned based on sample type (*i.e.,* IBC_LN samples vs. IBC_Pure samples), and 70% of samples were used as a training set, with the remaining 30% used as an independent test set. By comparing the IBC_LN- and IBC_Pure-derived sEV proteomes, we discovered that the best partition was achieved with 12 sEV proteins (PEPD, NCL, PARP1, ACTA2, ACTG2, TBCA, TTYH3, MATR3, KPNB1, KRT16, RANBP2, and CCT6A). Based on this 12-protein signature, applying 5-fold cross-validation to the training set yielded a sensitivity (true positive rate) of 100% and specificity (true negative rate) of 100% (Fig 4J and L, Appendix Fig S4J). When applying the protein signature for discriminating BC patients with or without LN metastasis to the independent test set samples, it had a sensitivity of 81% and a specificity of 81% (Fig 4M and N, Appendix Fig S4E). In addition, we used the CPTAC breast cancer dataset (n = 77) as an external validation test set and achieved 100% sensitivity and 100% specificity (Mertins *et al*, 2016) (Fig 4K).

### Potential sEV survival biomarkers for distant metastases of BC

To identify universal biomarkers associated with distant metastasis, we performed further analysis based on the proteomic profiles of 7 ductal carcinoma in situ (DCIS) samples and 21 distant metastasis (D-MET) (*e.g.,* M-Multiple (n = 5), M-Lung (n = 3), M-Liver (n = 4), M-Bone (n = 7), M-Chest wall (n = 1), and M-Soft tissue (n = 1)) samples in our cohort. Clustering and cluster-specific enrichment analyses of the upregulated proteins using DAVID (KEGG gene sets) pathway annotations clearly showed distinctive biological processes and pathways enriched in D-MET samples compared to DCIS samples (Fig 5A). Compared with DCIS samples, D-MET samples showed an upregulation of focal adhesion (*i.e.,* FLNA and vitronectin (VTN)) (Fisher’s exact test, *p* = 5.45e-03), metabolism-related pathways (*e.g.*, carbon metabolism (*i.e.,* PKM, G6PD, and TALDO1) (Fisher’s exact test, *p* = 5.42e-05), glycolysis/gluconeogenesis (*i.e.,* FBP1, LDHB, and PDHB) (Fisher’s exact test, *p* = 1.33e-02), fatty acid metabolism (i.e., ACACA, HSD17B12, and HACD3) (Fisher’s exact test, *p* = 1.83e-02)), and complement and coagulation cascades (*i.e.,* CPB2, alpha-1-antitrypsin (SERPINA1), CFH, C7, heparin cofactor 2 (SERPIND1), F10, F12, SERPINF2, SERPINE1, F2, TFPI, and KNG1) (Fisher’s exact test, *p* = 3.58e-02) (Fig 5A and B). We found that 24 sEV proteins were significantly overexpressed in distant metastatic samples (D-MET_median_/DCIS_median_ > 2-fold, Student’s *t* test, *p* < 0.05), suggesting that they may be potential serum sEV protein markers for LN metastasis of BC (Fig 5B). Among them, 5 sEV proteins (PDHB, FBP1, PPP4C, GP1BA, and TFPI) were identified in > 75% of D-MET samples (Fig 5B). Remarkably, 11 sEV proteins (FLNA, VTN, PKM, PDHB, G6PD, TALDO1, LDHB, ACACA, PPP4C, C7 and F2) were highly expressed in BC and were associated with poor prognosis in the Tang et al. BC cohort and the Liu et al. BC cohort (Liu *et al*, 2014; Tang *et al*., 2018) (Fig 5B and C, Appendix Fig S5A).

**Figure 5.**
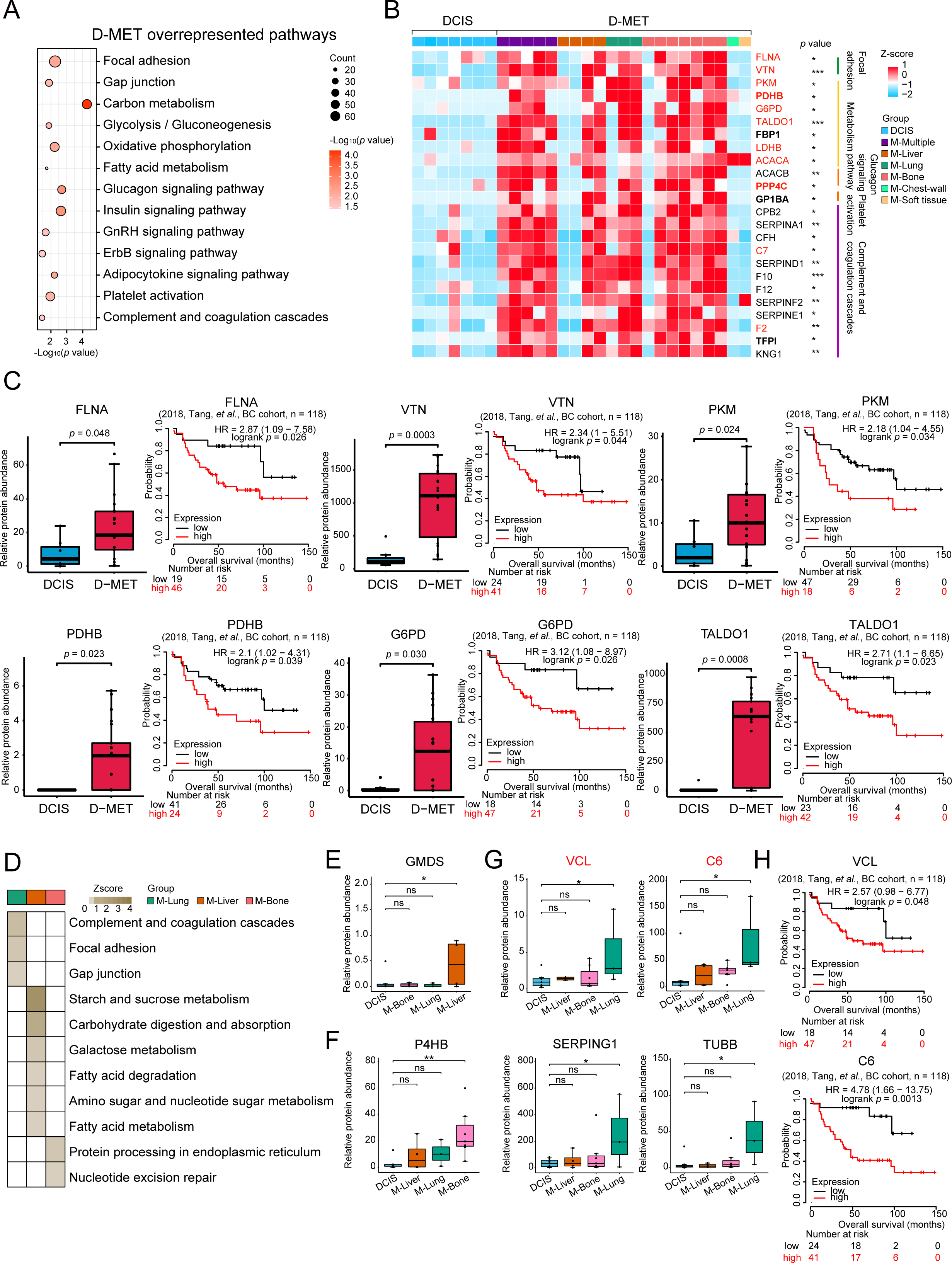
Potential sEV survival biomarkers for the distant metastases of BC. A The bubble plot indicates the overrepresented pathways in D-MET compared to DCIS. See Table S5. B Differentially expressed proteins between distant metastases and DCIS samples with > 2-fold difference and two-way Student’s *t* test *p* < 0.05. C Potential markers of distant metastasis were significantly associated with clinical outcomes in BC (2018, Tang et al., BC cohort, n = 118) (*p* value from log rank test). D DAVID (KEGG gene sets) analyses of the proteomic data of 21 BC patients with distant metastases revealed pathways that were significantly altered in lung metastases (M-Lung, n = 3), liver metastases (M-Liver, n = 4), and bone metastases (M-Bone, n = 7) (Fisher’s exact test, *p* < 0.05). E GMDS was specifically highly expressed in M-Liver. ns, no significance; \**p* < 0.05 by one-way Student’s *t* test. F P4HB was specifically highly expressed in M-Bone. ns, no significance; \*\**p* < 0.01 by one-way Student’s *t* test. G C6, TUBB, SERPING1 and VCL were specifically highly expressed in M-Lung. ns, no significance; \**p* < 0.05, \*\**p* < 0.01 by one-way Student’s *t* test. H High expression of VCL was associated with poor prognosis in BC (2018, Tang, et al. BC cohort, n = 126).

### Potential organ-specific sEV survival biomarkers for distant metastases of BC

Furthermore, we performed pathway enrichment analysis comparing differentially expressed proteins among three different types of organ metastasis samples (M-Lung, M-Liver, and M-Bone samples). M-Lung sEVs showed upregulation of complement and coagulation cascades (*i.e.,* CFD, C6, and SERPING1) (Fisher’s exact test, *p* = 1.36e-02), focal adhesion (*i.e.,* ITGB3, ITGA2B, and VCL) (Fisher’s exact test, *p* = 2.63e-02), and gap junctions (*i.e.,* TUBB2B, TUBB2A, and TUBB) (Fisher’s exact test, *p* = 3.49e-02) (Fig 5D, Appendix Table 5). This finding is consistent with recent reports that focal adhesion and regulation of actin cytoskeleton signaling are involved in lung metastases of BC (Zeng *et al*, 2019). Interestingly, we found that abundant metabolism-related pathways were enriched in M-Liver sEVs, including fatty acid metabolism (*i.e.,* ACADVL, TECR, and ACSL5) (Fisher’s exact test, *p* = 4.56e-02), galactose metabolism (*i.e.,* GLB1, PGM5, and PGM1) (Fisher’s exact test, *p* = 1.90e-02), and starch and sucrose metabolism (*i.e.,* AMY2A, AMY1A, and AMY2B) (Fisher’s exact test, *p* = 7.99e-05) (Fig 5D, Appendix Table 5). M-Bone sEV samples showed upregulation of protein processing in the endoplasmic reticulum (*i.e.,* HSPH1, STT3A, RAD23A, P4HB, and SEC23B) (Fisher’s exact test, *p* = 1.90e-02) and nucleotide excision repair (*i.e.,* RPA1, RAD23A, and CUL4B) (Fisher’s exact test, *p* = 3.31e-02) (Fig 5D, Appendix Table 5). These results suggest that although upregulated expression of adhesion, metabolism, and angiogenesis pathways are common features of distant metastases, different metastases are biased. M-Lung was the adhesion type, M-Liver was the metabolism type, and M-Bone was the repair type.

We found that GMDS was specifically highly expressed in M-Liver, P4HB was specifically highly expressed in M-Bone, and C6, TUBB, SERPING1 and VCL were specifically highly expressed in M-Lung (Fig 5E-G). In the Tang et al. BC cohort, the high expression of C6 and VCL was associated with poor prognosis, suggesting that they may be survival markers for lung metastasis of BC (Tang *et al*., 2018) (Fig 5G and H, Appendix Fig S5B).

### Potential BC-derived sEV molecules govern organ-specific metastasis

Metastatic organotropism has remained an enigmatic issue. A recent study showed that cancer-derived sEV uptake by organ-specific cells may govern organ-specific metastasis (Hoshino *et al*., 2015). To examine whether sEV proteins may guide the colonization of BC cells in specific organs, we computed the abundance of specific cell types in each of the distant metastatic samples using xCell (Fig 6 A-D, Appendix Fig S6A and B, Appendix Table 6). The analysis showed an enhanced enrichment score of chondrocytes in M-Bone sEVs, which was 6-fold, 1.75-fold, and 2.75-fold higher than that in DCIS, M-Lung, and M-Liver sEVs, respectively (Fig 6B). In contrast, the enrichment score of myocytes in M-Lung sEVs was upregulated by 2.20-fold, 5.32-fold, and 1.82-fold compared to that in DCIS, M-Liver, and M-Bone sEVs, respectively (Fig 6C). Moreover, the enrichment score of fibroblasts in M-Liver samples was significantly elevated by 4.10-fold, 12.77-fold, and 5.33-fold compared to DCIS, M-Lung, and M-Bone samples, respectively (Fig 6D) (Student’s *t* test, *p* < 0.05). Therefore, the organ specificity of sEV biodistribution matched the organotropic distribution of tumor cells.

**Figure 6.**
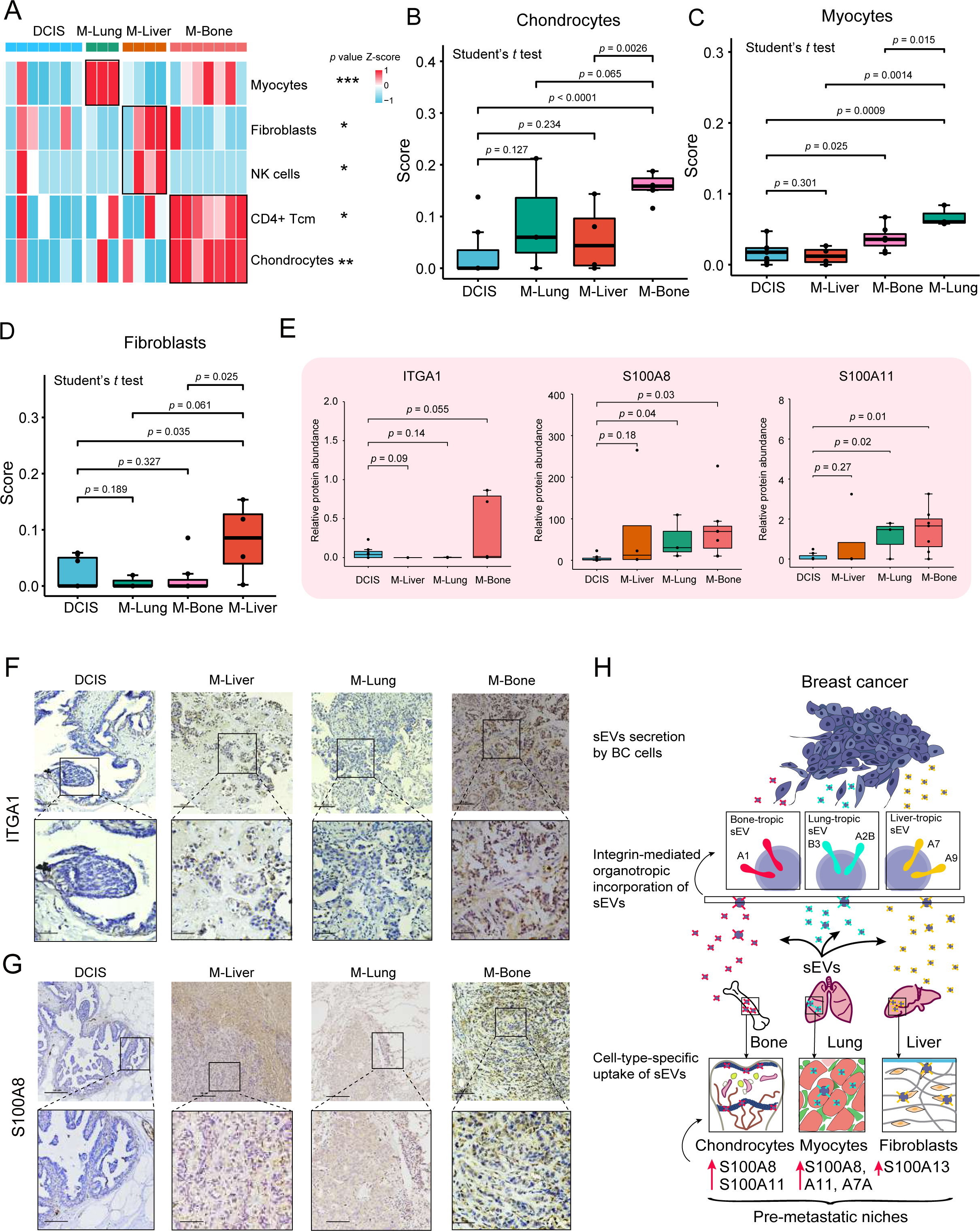
Potential BC-derived sEV molecules govern organ-specific metastasis. A Distinctive tumour microenvironment of M-Lung, M-Liver, and M-Bone samples. \**p* < 0.05, \*\**p* < 0.01, \*\*\**p* < 0.001 by ANOVA. B Boxplot showing the relative abundance of chondrocytes in the distant metastases of BC. *P* value from one-way Student’s *t* test. C Boxplot showing the relative abundance of myocytes in the distant metastases of BC. *P* value from one-way Student’s *t* test. D Boxplot showing the relative abundance of fibroblasts in the distant metastases of BC. *P* value from one-way Student’s *t* test. E sEV ITGA1, S100A8 and S100A11 molecular levels in M-Bone. *P* value from one-way Student’s *t* test. F Protein expression of ITGA1 in DCIS, M-Liver, M-Lung, and M-Bone tissues detected by using immunohistochemistry. G Protein expression of S100A8 in DCIS, M-Liver, M-Lung, and M-Bone tissues detected by using immunohistochemistry. H Model of sEV-mediated organotropic tumour dissemination. BC-derived sEVs are taken up by organ-specific resident cells in metastatic organs based on integrin expression.

A previous study suggested that specific exosomal integrins were associated with metastatic organotropism by dictating premetastatic niche formation (Hoshino *et al*., 2015). In our dataset, we identified 25 integrins enriched in M-Bone, M-Lung and M-Liver sEVs. Further analysis revealed that ITGA1 was primarily detected in M-Bone sEVs, ITGA7 and ITGA9 were abundantly enriched in M-Liver sEVs, and ITGB3 and ITGA2B were abundantly enriched in M-Lung sEVs (Fig 6E, Appendix Fig S6C and D).

In addition to adhesive properties, sEV integrins can upregulate promigratory and proinflammatory S100 molecules, which influence premetastatic niche formation (Hoshino *et al*., 2015). To determine the pattern of sEV-S100 molecules in tumor metastasis, we identified 16 S100 molecules from M-Bone, M-Lung and M-Liver sEVs. The analysis revealed that S100A8 was primarily detected in sEV-derived proteins from M-Bone samples (Fig 6E). S100A13 was primarily detected in M-Liver samples (Appendix Fig S6C). Interestingly, S100A7A was abundantly present in sEV-derived proteins from M-Lung samples (Appendix Fig S6D). In addition, we verified that ITGA1 was significantly increased in M-Bone tissue samples in our additional BC cohort (DCIS (n = 4), M-Liver (n = 4), M-Lung (n = 4), and M-Bone (n = 8)) (Student’s t test, *p* < 0.05) (Fig 6F, Appendix Fig S6A). Consistently, S100A8, S100A13, and S100A7A were significantly increased in M-Bone, M-Liver, and M-Lung tissue samples in our additional BC cohort (DCIS (n = 3), M-Liver (n = 3), M-Lung (n = 4), and M-Bone (n = 8)) (Student’s *t* test, *p* < 0.05) (Fig 6G, Appendix Fig S6B, E and F). Taken together, these results suggested a correlation between specific sEV integrins and S100 molecules and tissue organotropism (Fig 6H).

## Discussion

Blood tests remain the most readily accessible source for the early detection, classification, and treatment guidance of BC patients. The billions of sEVs circulating in blood could represent an essential component of liquid biopsy (Miyagi *et al*, 2021). Despite previous studies on BC-derived sEVs (Chen *et al*, 2017), there is a lack of a comprehensive understanding of BC-specific sEV characteristics and their composition and consensus on unique BC biomarkers due to limited sEV proteome data from human samples.

Here, we performed a large-scale comprehensive analysis of sEV proteomes from 167 serum samples obtained from patients with BC, patients with BD, and healthy individuals. Firstly, we applied this eight-protein (STAT1, PON1, APOC1, APOC2, MMP2, IGHV4-39, IGHV3-53, and ADD2) identifier to sEV samples of the independent test set, the model achieved 97% sensitivity and 83% specificity in the diagnosis of BC. This study may provide reference value for differentiating benign and malignant breast tumors using serum in the future.

BC is a heterogeneous disease in terms of molecular alterations, cellular compositions, and clinical outcomes (Wagner *et al*, 2019). Therefore, the classification of molecular subtype is an important tool for treatment and prognosis evaluation. Clinically, based on the expression of ER, PR, Her2, and Ki67 by IHC, BC is categorized into various molecular subtypes (Holm *et al*, 2021). However, the patterns of these biological indicators may change during the course of BC progression, so they may be used to adjust treatment strategies accordingly (Ju *et al*, 2018). Thus, we speculated that an sEV-based in vitro diagnostic strategy is an emerging approach complementary to tissue pathology. Unfortunately, we failed to confirm the existence of ER, PR, HER2, and Ki67 in the serum sEV datasets, indicating that they may either have a low abundance or be lacking in serum sEVs. However, further analyses of differentially regulated sEV-derived proteins in luminal A, luminal B, Her2-enriched, and TNBC samples clearly showed significant differences in the proteins and biological pathways involved. By comparing proteomic profiles among diverse molecular subtypes of BC, we constructed a 61-protein classifier. The ROC curve derived from the 61-protein signature showed good sensitivity and specificity, with an AUC of 1.0. Then, the 61-protein signature was validated in the test set, resulting in a ROC curve with an AUC of 0.875. This work may provide reference value for the diagnosis of clinical subtypes of BC using serum in the future.

An accurate preoperative assessment of LN status is one of the most important prognostic factors determining the long-term outcome (Banerjee *et al*, 2004). Although noninvasive imaging modalities such as ultrasonography, computed tomography, and magnetic resonance imaging have been widely adopted for the clinical evaluation of LN status before surgery, the sensitivity of these modalities is not satisfactory (Song, 2020). In the present study, PCA demonstrated a clear distinction between IBC_LN samples and IBC_Pure samples, which further highlighted the diverse proteomic patterns between IBC with or without LN metastasis. Hence, we constructed an sEV-based protein signature that predicted LN metastasis at the serum sEV proteomic level based on machine learning classification, showing 81% and 81% specificity and sensitivity, respectively. In addition, we used the CPTAC BC dataset (n = 77) as an external validation test set and achieved 100% sensitivity and 100% specificity. These data suggest that tumor-associated sEV proteins can serve as biomarkers for early-stage cancer detection of LN metastasis.

Previous studies showed that adhesion and ECM molecules, such as integrins, tenascin and periostin, were associated with distant metastasis of disseminating cancer cells (Fukuda *et al*, 2015; Oskarsson *et al*, 2011; Radisky *et al*, 2002; Weaver *et al*, 1997). Regarding the research on this aspect, Hoshino et al. defined a specific repertoire of integrins expressed on cancer-derived exosomes, distinct from cancer cells, that dictate metastatic tropism (Hoshino *et al*., 2015). In our study, we identified 25 integrins abundantly present in human bone-, lung- and liver-tropic metastatic sEVs by quantitative mass spectrometry. Notably, we found that sEVs expressing ITGA1 may specifically bind to chondrocytes, which are related to bone tropism. sEVs expressing ITGB3 and ITGA2B may specifically bind to lung-resident myocytes, mediating lung tropism. However, sEVs expressing ITGA7 and ITGA9 may bind liver-resident fibroblasts, governing liver tropism. Moreover, we revealed that the pattern of sEV-S100 molecules was correlated with tissue organotropism and could serve as a biomarker for distant metastasis (Fig 6H).

In conclusion, our findings show that proteins carried by BC-derived sEVs could be used as a novel, minimally invasive liquid biopsy tool for the early detection of BC, as well as for discriminating molecular subtypes, LN involvement status, and organotropic metastasis. These findings could advance the implementation of routine serum sEV-based screening in the clinic.

## Materials and Methods

### Sample collection

Serum sample collection was approved by Shanghai General Hospital Shanghai Jiao Tong University School of Medicine (Shanghai, China, permit number [2017]KY053), and all patients provided proper consent before samples were collected. Serum samples were collected between March 2011 and August 2019. Detailed information is shown in Appendix Table 1.

### sEV extraction

Isolation of exosomes was performed by differential ultracentrifugation following established centrifugation times and parameters (An *et al*, 2018; Gao *et al*, 2021; Lakhter *et al*, 2018; Takov *et al*, 2019; Thery *et al*, 2006). Firstly, 1 mL serum was thawed on ice and centrifuged at 3,000 g for 10 min at 4°C. The supernatant was removed, and large vesicles were removed with another centrifugation step at 10,000 g for 20 min at 4°C and the supernatant was diluted with 25 mL PBS and filtered through a 0.22 µm centrifugal filter device to remove any large contaminating vesicles. Secondly, filtered serum was centrifuge at an overspeed of 150,000 g for 4 h, the milky white floating object at the top was sucked away. Thirdly, centrifuged material was resuspended with 25 mL PBS and further centrifuged at 4°C for 150,000 g for 2 h. Fourthly, supernatant was discarded and 200 µL solution was retained at the bottom to resuspend the precipitate. Isolation and relative purity of the sEVs were confirmed by NTA, transmission electron microscopy (TEM) and immunoblot.

### sEVs protein extraction and tryptic digestion

sEV samples (typically 5 µg, adjusted based on BCA measurements) were dried by vacuum centrifugation and redissolved in 30–50 µL of 8 M urea/50 mM ammonium bicarbonate/10 mm DTT. Following lysis and reduction, proteins were alkylated using 20 or 30 mM iodoacetamide (Sigma, St. Louis, MO, USA). Proteins were digested with trypsin (Promega, Madison, WI, USA) at an enzyme-to-protein mass ratio of 1:50 overnight at 37°C, and peptides were then extracted and dried (SpeedVac, Eppendorf). Peptides were desalted and concentrated using Empore C_18_-based solid phase extraction prior to analysis by high resolution/high mass accuracy reversed-phase (C_18_) nano-LC-MS/MS.

### Liquid chromatography

We employed an EASY-nLC 1200 ultra-high-pressure system liquid chromatography system (Thermo Fisher Scientific). Peptides were separated within 75 min at a flow rate of 600 nL/min on a 150 μm I.D. × 15 cm column with a laser-pulled electrospray emitter packed with 1.9 μm ReproSil-Pur 120 C_18_-AQ particles (Dr. Maisch). Mobile phases A and B were water and acetonitrile with 0.1 vol% FA, respectively. The %B was linearly increased from 15 to 30% within 75 min.

### Mass spectrometry

Samples were analysed on a Q-Exactive-HF mass spectrometer (Thermo Fisher Scientific) via a nanoelectrospray ion source (Thermo Fisher Scientific). The mass spectrometer was operated in data-independent mode for ion mobility-enhanced spectral library generation. Typically, 75% of samples were injected. The peptides were dissolved in 12 μL of loading buffer (0.1% formic acid), and 9 μL was loaded onto a 100 μm I.D. × 2.5 cm C_18_ trap column at a maximum pressure of 280 bar with 14 μL of solvent A (0.1% formic acid). The DIA method consisted of an MS1 scan from 300–1400 m/z at 60 k resolution (AGC target 4e5 or 50 ms). Then, 30 DIA segments were acquired at 15 k resolution with an AGC target of 5e4 or 22 ms for maximal injection time. The setting “injections for all available parallelizable time” was enabled. HCD fragmentation was set to a normalized collision energy of 30%. The spectra were recorded in profile mode. The default charge state for the MS2 scan was set to 3.

### Peptide identification and protein quantification

All data were processed using Firmiana (Feng *et al*, 2017). The DIA data were searched against the UniProt human protein database using FragPipe (v.12.1) with MSFragger (2.2) (Kong *et al*, 2017). The mass tolerances were 20 ppm for precursor and 50 mmu for product ions. Up to two missed cleavages were allowed. The search engine set cysteine carbamidomethylation as a fixed modification and N-acetylation and oxidation of methionine as variable modifications. Precursor ion score charges were limited to +2, +3, and +4. The data were also searched against a decoy database so that protein identifications were accepted at a false discovery rate (FDR) of 5%.

The DIA data was analysed using DIANN (v.1.7.0) (Demichev *et al*, 2020). The quantification of identified peptides was calculated as the average chromatographic fragment ion peak areas across all reference spectra libraries. Label-free protein quantifications were calculated using a label-free, intensity-based absolute quantification (iBAQ) approach (Zhang *et al*, 2012). We calculated the peak area values as parts of the corresponding proteins. The fraction of total (FOT) was used to represent the normalized abundance of a particular protein across samples. FOT was defined as a protein’s iBAQ divided by the total iBAQ of all identified proteins within a sample. The FOT values were multiplied by 10^5^ for ease of presentation, and missing values were imputed to 10^-5^. The raw proteomics data files are hosted by iProX and can be accessed at https://www.iprox.cn (Project ID: IPX0003429000).

### Statistical analysis

To impute the proteomic data, we first screened more than 50% of the identified proteins in each group and divided the data into two parts. When the protein detection rate was < 50%, the missing value was replaced with one tenth of the minimum value. For these proteins, no imputation was applied. When the protein detection rate was > 0.5, the missing value was probably due to the detection accuracy limitation of LC/MS. In this case, we first calculated the missing probability of a protein using the R package “impute” (https://git.bioconductor.org/packages/impute) based on the K-NN algorithm. Meta-analysis-based discovery and validation of survival biomarkers was carried out using Kaplan-Meier Plotter (http://kmplot.com/analysis/).

### Principal component analysis (PCA)

The imputed data were then normalized using the LogNorm algorithm. The PCA function of the R package “factoextra” (https://cran.r-project.org/web/packages/factoextra/index.html) was used to implement unsupervised clustering analysis. The 95% confidence coverage was represented by a coloured ellipse for each group and was calculated based on the mean and covariance of points in the different groups. 1,734, 1,038, and 1,116 proteins (features) were used for PCA to illustrate the global proteomic differences among BC (n = 126), BD (n = 17) and HD (n = 24), the global proteomic differences among the luminal A (n = 20), luminal B (n = 50), Her2-enriched (n = 21), and triple-negative breast cancer (TNBC) (n = 23) subtypes, and the global proteomic differences between IBC_Pure (n = 54) and IBC_LN (n = 51) (Appendix Fig S2A, 3A, 4A).

### Global Heatmap

Each gene expression value in the global proteomic expression matrix was transformed to a z-score across all the samples. The z-score-transformed matrix was clustered using the R package “pheatmap” (https://cran.r-project.org/web/packages/pheatmap/index.html).

### Pathway enrichment analysis

Pathway enrichment analysis was performed by DAVID (https://david.ncifcrf.gov) and ConsensusPathDB (http://cpdb.molgen.mpg.de), and significance in the pathway enrichment analysis was determined by Fisher’s exact test on the basis of Kyoto Encyclopedia of Genes and Genomes (KEGG) pathways and categorical annotations, including Gene Ontology (GO) biological process (GOBP) terms and Reactome (https://reactome.org).

### Multiplex immunohistochemistry (mIHC) with tyramide signal amplification

Tissues or cells were prepared for detection with kits using standard fixation and embedding techniques. Each slide was baked in an oven at 65°C for 1 h, dewaxed with xylene (3 x 10 min) and rehydrated through a graded series of ethanol solutions (100% ethanol, 95% ethanol, 75% ethanol, 50% ethanol)and each step took 5 min. After rehydration, immersing the slides in the boiled appropriate AR buffer, and placed in a microwave for 15 min at 20% power. After naturally cooling to room temperature, washing the slides with TBST. Then we used blocking buffer to incubate tissue section for 10 min. The blocking buffer was drained, and Primary Antibody Working Solution was applied. CD45RA (1:3000; ab755; Abcam), CD34 (1:6000; ab81289; Abcam), CD38 (1:800; ab108403; Abcam), CD71 (1:800; ab214039; Abcam), and CDH1 (1:10000; ab181860; Abcam) were used. The slides were incubated at 4°C overnight or at room temperature for 1 h; the time may be adjusted according to different characteristics of the antibody. After washing the slides with TBST, incubate them in polymer HRP Ms+Rb for 15 min at room temperature. Washing the slides twice again. Working Solution (100–300 µL) was pipetted onto each slide at room temperature for 10 min. And then immersed in the appropriate AR buffer. This microwave step strips the primary-secondary-HRP complex, allowing the introduction of the next primary antibody. For detection of the next target with fluorophores, we restarted the protocol at blocking. Once all 5 targets were labelled, Opal Polaris 780 labelling was continued.

Dropping TSA-DIG Working Solution onto slides and incubating at room temperature for 10 min. Repeat the previous microwave repair steps after washing the slides. Polaris 780 Working Solution was pipetted onto each slide and incubated at room temperature for 1 h. DAPI working solution was applied for 5 min. The slides were washed twice again. After the slides were slightly dry, a super quench sealing tablet was added to the slides with a pipette, and the sample area was immersed.

### Immunohistochemistry (IHC)

Firstly, the sections were baked at 65°C for 1 h and incubated in xylene three times for 10 min each time. Then, the sections were hydrated by a graded series of ethanol (100% ethanol, 95% ethanol, 75% ethanol, 50% ethanol and ddH_2_O), and each step took 5 min. Antigen retrieval was conducted using a microwave oven: 3 min at 100% power and 15 min at 20% power filled with 10 mM sodium citrate buffer (pH 6.0). After naturally cooling to room temperature and washed in ddH_2_O, we blocked the sections with 5% normal goat serum for 10 min, incubated sections in 3% H_2_O_2_ for 10 min at room temperature, and washed the sections twice in PBS for 5 min. The following antibodies were diluted in the appropriate concentrations: PPARg (1:10000; ab59256; Abcam), S100A8 (1:800; 15792–1-AP; Proteintech), S100A13 (1:1200; ab109252; Abcam), S100A7A (1:400; DF8517; Affinity), and ITGA1 (1:300; 22146–1 AP; Proteintech). These antibodies were incubated with the sections overnight at 4°C.

The next day, after washing the sections twice in PBS, we used an IHC Kit (ZSGB-BIO, Beijing, China, Cat# SP-9000), incubated the sections with biotin-labelled secondary antibody for 15 min. After washing sections twice in PBS, incubating the sections with horseradish enzyme-labelled Streptomyces ovalbumin working solution for 15 min. Finally, We used DAB solution to stain the tissues. Then, using haematoxylin to stain nuclears and washing them in ddH_2_O. Finally, the sections were dehydrated by graded ethanol (50% ethanol, 75% ethanol, 95% ethanol, and 100% ethanol). We dried the slides in a fume cupboard for at least 20 min and mounted coverslips.

## Acknowledgments

This work was supported by the National Natural Science Funds (grant numbers 82073269, 81772802 and M-0349), Shanghai Science and Technology Innovation Action Plan (grant number 20XD1402800), Clinical Research Plan of SHDC (grant number SHDC2020CR2065B), Clinical Research Innovation Plan of Shanghai General Hospital (grant number CTCCR-2016B05), National Key R&D Program of China (grant numbers 2017YFA0505102, 2016YFA0502500, 2018YFA0507501, and 2017YFC0908404), National Natural Science Foundation of China (grant numbers 31770886, 1972933, and 31700682), Science and Technology Commission of Shanghai Municipality (grant number 2017SHZDZX01), Major Project of Special Development Funds of Zhangjiang National Independent Innovation Demonstration Zone (grant number ZJ2019-ZD-004).

## Author Contributions

H.X.W. contributed to idea, conception, and study design. H.X.W., C.D., and H.W.Z. wrote the paper and supervised the project. G.F.X., S.Q.D. and M.J.H. conducted the mass spectrometry analysis. W.Y.H. and the other authors carried out all the remaining experiments. All authors discussed the results, commented on the project and approved the manuscript.

**Appendix Figure 1. Proteomic characterization of BC-derived sEVs, related to Figure 1**

A NanoSight profiles showing the size distribution of serum-derived sEVs isolated from BC, BD, and HD. Red denotes BC-derived sEVs, yellow denotes BD-derived sEVs, and blue denotes HD-derived sEVs.

B Identification of 24 sEV protein markers in our proteomic data.

C Distribution of log10-transformed iBAQ abundance of identified proteins in 167 proteome samples that passed quality control. Red denotes BC samples (n = 126), yellow denotes BD samples (n = 17), and blue denotes HD samples (n = 24). In the box plots, the middle bar represents the median, and the box represents the interquartile range; bars extend to 1.5 × the interquartile range.

**Appendix Figure 2. Proteomics features of BC-, BD- and HD-derived sEVs, related to Figure 2**

A PCA of 1,734 proteins in 167 samples. Red, BC (n = 126); yellow, BD (n = 17); blue, HD (n = 24).

B Schematic diagram of the structural distribution of damage-associated molecular patterns (DAMPs) in sEVs (left). Venn diagram showing the number of DAMPs detected in BC, BD, and HD samples (right). See Table S2.

C sEV DAMP molecules enriched in BC were significantly associated with clinical outcomes in BC (2018, Tang et al., BC cohort, n = 118) (*p* value from log rank test).

D The dataset was split randomly into training (70%) and test sets (30%) at the patient level. A machine learning algorithm, XGBoost, was used for model development, training, and validation. Receiver operating characteristic (ROC) analysis was used to evaluate the performance of the classifier on the test dataset.

**Appendix Figure 3. Proteomic landscapes of four clinical subtypes of BC-derived sEVs, related to Figure 3**

A PCA of 1,308 proteins in 114 samples. Orange, luminal A (n = 20); green, luminal B (n = 50); purple, Her2-enriched (n = 21); and blue, TNBC (n = 23).

B Proteins with the highest predictive values in classifying luminal A, luminal B, Her2-enriched, and TNBC samples by XGBoost.

**Appendix Figure 4. Potential prognostic biomarkers for IBC patients with lymph node metastases, related to Figure 4**

A PCA of 1,116 proteins in 105 samples. Blue, invasive breast cancer with lymph node metastases (IBC_Pure, n = 54); red, invasive breast cancer without lymph node metastases (IBC_LN, n = 51).

B Differentially expressed proteins between IBC_Pure and IBC_LN samples that were found in > 50% of the corresponding samples, with > 2-fold difference and Student’s *t* test *p* < 0.05.

C Comparison of the scores of adipocytes between the IBC_LN group and the IBC_Pure group. The *p* value was calculated by the Wilcoxon rank sum test. The line and box represent median and upper and lower quartiles, respectively.

D Correlation between adipogenesis and adipocytes. Spearman rho = 0.188, *p* value = 5.507e-02.

E Comparison of the MPP scores between the IBC_LN group and the IBC_Pure group. The *p* value was calculated by the Wilcoxon rank sum test. The line and box represent the median and upper and lower quartiles, respectively.

F Correlation between MPPs and the coagulation pathway. Spearman rho = 0.295, *p* value = 2.216e-03.

G Correlation between platelets and the coagulation pathway. Spearman rho = 0.209, *p* value = 3.225e-02.

H Comparison of the platelet scores between the IBC_LN group and the IBC_Pure group. The *p* value was calculated by the Wilcoxon rank sum test. The line and box represent the median and upper and lower quartiles, respectively.

I Molecules that are highly associated with platelets.

J Representative fluorescence microscopy images of MEPs labelled with CD71 (green), CD38 (red), and CD45RA (yellow). Images revealed the presence of MEPs in lymph node metastases of BC, which were rare in normal lymph nodes and primary breast cancer.

K Platelet counts in the blood of IBC patients with lymph node metastasis (n = 43) and IBC patients without lymph node metastasis (n = 45).

L The dataset was randomly split into training (70%) and test sets (30%) at the patient level. A machine learning algorithm, XGBoost, was used for model development, training, and validation. Receiver operating characteristic (ROC) analysis was used to evaluate the performance of the classification on the test dataset.

**Appendix Figure 5. Potential sEV survival biomarkers for the distant metastases of BC, related to Figure 5**

A Potential markers of distant metastasis were significantly associated with clinical outcomes in BC (2018, Tang et al., BC cohort, n = 118 and 2014, Liu et al., BC cohort, n = 126) (*p* value from log rank test).

**Appendix Figure 6. Potential molecules present on IBC-derived sEVs target them to specific organs, related to Figure 6**

A IHC score of ITGA1 in DCIS (n = 4), M-Liver (n = 4), M-Lung (n = 4), and M-Bone (n = 8). *P* value from two-way Student’s *t* test.

B IHC score of S100A8 in DCIS (n = 3), M-Liver (n = 3), M-Lung (n = 4), and M-Bone (n = 8). *P* value from two-way Student’s *t* test.

C sEV ITGA7, S100A9 and S100A13 molecular levels in M-Liver. *P* value from one-way Student’s *t* test.

D sEV ITGB3, S100A2B and S100A7A molecular levels in M-Lung. *P* value from one-way Student’s *t* test.

E The protein expression of S100A13 in DCIS (n = 3), M-Liver (n = 3), M-Lung (n = 4), and M-Bone (n = 8) tissues was detected by using immunohistochemistry (left); IHC score of S100A13 in M-Liver. *P* value from two-way Student’s *t* test (right).

F The protein expression of S100A7A in DCIS (n = 3), M-Liver (n = 3), M-Lung (n = 4), and M-Bone (n = 8) tissues was detected by using immunohistochemistry (left); IHC score of S100A7A in M-Lung. *P* value from two-way Student’s *t* test (right).

